# Unsupervised deep learning method for cell segmentation

**DOI:** 10.1101/2021.05.17.444529

**Authors:** Nizam Ud Din, Ji Yu

## Abstract

Advances in the artificial neural network have made machine learning techniques increasingly more important in image analysis tasks. Recently, convolutional neural networks (CNN) have been applied to the problem of cell segmentation from microscopy images. However, previous methods used a supervised training paradigm in order to create an accurate segmentation model. This strategy requires a large amount of manually labeled cellular images, in which accurate segmentations at pixel level were produced by human operators. Generating training data is expensive and a major hindrance in the wider adoption of machine learning based methods for cell segmentation. Here we present an alternative strategy that uses unsupervised learning to train CNNs without any human-labeled data. We show that our method is able to produce accurate segmentation models. More importantly, the algorithm is applicable to both fluorescence and bright-field images, requiring no prior knowledge of signal characteristics and requires no tuning of parameters.

## Introduction

Automated cellular segmentation from optical microscopy images is a critical task in many biological researches that rely on single-cell analysis. Traditional approaches to cell segmentation rely on manually-crafted feature definitions that allow the algorithmic recognition of cellular area and cell border^1,2^. Unfortunately manual feature definitions are usually highly context-specific and require task-dependent tuning to work well. For example, an algorithm that is well-optimized for a specific membrane fluorescent marker will not work on images of a different fluorescent marker, nor on bright field images. Switching to a different cell type (with different morphological features), changing imaging modality (e.g., from epi-fluorescence to confocal), or even imaging settings (e.g. changing the objective) often requires a redesign/reoptimization of the segmentation algorithm. Therefore, there is a need for more generic, “turn-key” solutions for this task.

Convolutional neural networks (CNN) have recently been shown to be highly efficient on various kinds of image processing tasks, including semantic segmentation^3–6^. In particular, various networks with an hourglass-shaped architecture (e.g., UNet) have achieved remarkable pixel-level accuracy in segmentation of objects, including biological cells^7–11^. However, there are two major problems that have prevented the adoption of CNNs in biological research. First, while CNNs perform well in segmentation of a single cell, they often exhibit an under-segmentation bias when processing crowded cell populations that contain many cells adjacent to each other^8^. This makes them less desirable for common analytical tasks such as high content screening, in which typically hundreds to thousands of cells need to be segmented simultaneously from one image dataset. Second, and more importantly, to train a CNN for the segmentation task, one needs a significant amount of manually labeled training images, in which cell areas and/or cell boundaries are marked by human operators. Manually labeling is a time-consuming and expensive process. As a result, CNN-based approach remains uncompetitive versus traditional segmentation approaches in real-life biological applications.

Here we propose a new CNN-based cell segmentation method that aims to solve the two problems described above. A key difference between our method and previous attempts at the CNN-based cell segmentation is that we use a maker-controlled segmentation algorithm, taking a note from conventional cell segmentation pipelines. Many conventional segmentation algorithms (e.g., watershed^12^) are known to have an over-segmentation bias when the images are contaminated with noise. One way to correct for the bias is to provide a set of specific marker positions, depicting the approximate locations of each cell. Indeed, it is a common practice to use two-channel imaging data for cell segmentation, one channel for imaging nucleus and the other for the whole cell. The nucleus images are useful for computing marker positions, and the whole cell images provide information for determining cell boundaries. This approach is used in some of the most widely-used cell segmentation softwares, such as Cell Profiler^13^ and Ilastik^14^. The success of this approach derives partly from the fact that nucleus images have simple morphological features and therefore are relatively easy to analyze with traditional image processing algorithms. Nucleus identification and/or segmentation is a heavily researched topic; multiple algorithms exist in the literature with good performances^15–19^. The main challenge in cell segmentation is to devise reliable feature representations that can recognize cell boundaries with a high accuracy.

We propose to adopt the marker-controlled segmentation approach, but apply it to the CNN-based segmentation. We will first generate marker locations from nucleus images, but use a CNN to model cell features and produce accurate segmentation at the whole cell level. Unlike previous works on CNN segmentation, which use the networks to process the whole imaging field of view, we designed our network to perform segmentation on a smaller patch of the input image centered around the marker positions (Fig.1). This converts the multi-cell segmentation problem to multiple single-cell segmentation problems, which in turn removes the undersegmention bias as long as the nuclei markers were correctly computed. In addition, it is important to note that the “nucleus image” does not have to be one that is experimentally acquired. Instead, we will also demonstrate an alternative approach, in which we generate synthetic “nucleus images” from the normal whole cell images, by using a pretrained CNN model. This technique is similar to the method first demonstrated by Ounkomol et. al^20^, in which they showed that CNNs can be trained to map one image modality (e.g. bright-field image) to a different one (e.g, fluorescence images of plasma membrane, nucleus etc).

A more important consequence of the marker-controlled segmentation approach is that it allows us to devise a method to train segmentation CNNs (Fig. 1) in an unsupervised fashion, and thus avoiding the manual labeling of the training dataset. The key for achieving this is to design a custom objective function:

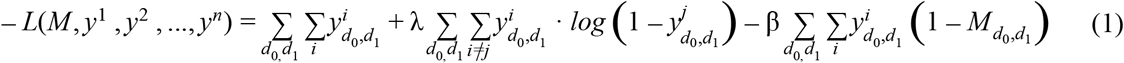

**Figure 1.**
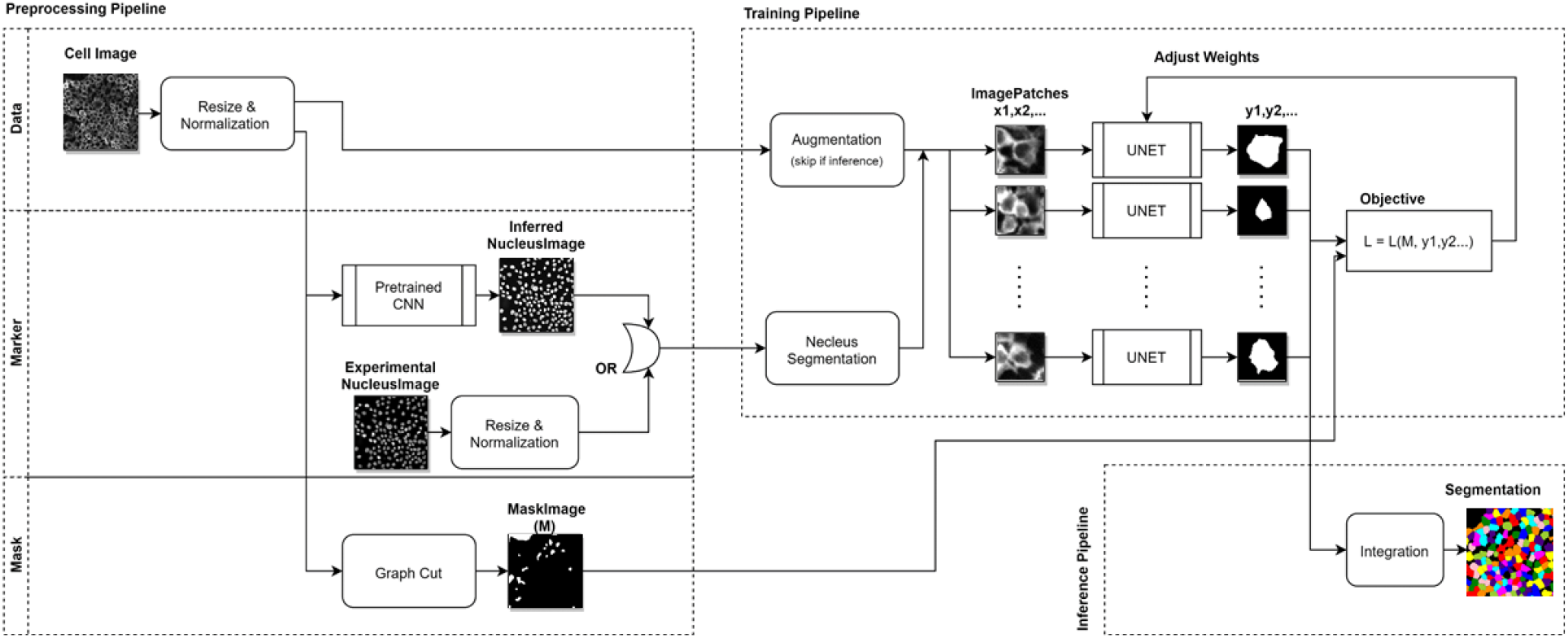
Schematic outline of the CNN segmentation algorithm. The overall process comprises three components: the preprocessing pipeline, which computes the marker location and image mask from the inputs, the training pipeline which extract image patches to train a CNN model for segmentation, and the integration pipeline that takes the single patch output from the CNN and create whole-field segmentation results.

Here, *y^i^* ∈ *R^k^*^×*k*^ is the CNN output from *i*-th patch of the input image, which can be interpreted as the probability value for classifying each pixel within the patch to be one belonging to the cell. The summations over *d*_0_,*d*_1_ are over all pixels in the input image, and the summations over *i* or *j* are over all patches. Furthermore, we use the notation 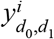 to denote the value of *y^i^* at a specific pixel location *d*_0_,*d*_1_. Since the individual CNN output covers only a small area of the entire image input, we assume a zero value when the index location *d*_0_,*d*_1_ is outside the computed area, consistent with the probabilistic interpretation of *y^i^*. Finally, *M* is a binary mask indicating the area of all cells, which can be generated from the input data with a graph cut algorithm. The goal is to train CNN until the loss function *L* is minimized. Since the loss function is designed to be fully differentiable, we can use the standard stochastic optimization^21^ and back-propagation technique to achieve the goal in a manner similar to supervised training.

The objective function contains three terms. The first term is optimized by including as many pixels as possible in the segmented results. This is balanced by the second term, which penalizes any overlaps of segmented pixels from different patches. Finally, the third term gives additional penalties if the segmented pixels exceed the masked area, i.e, it prevents the model from including the background pixels in the segmentation. The two hyper parameters, λ and β, control the relative weight of the penalty terms. Because the CNN model processes each individual patch independently, when segmenting one cell, the network is unaware of the results from adjacent patches. Therefore, the loss function is minimized only after the network has “learned” meaningful semantic features from the images, which in turn would hopefully result in correct segmentations.

## Results

We implemented the proposed algorithm using the TensorFlow computational framework^22^ and tested its efficacy on both immunofluorescence (anti-phosphortyrosine (pY)) and bright-field microscopy image data. In addition, to compute marker locations, we obtained nucleus images by staining cells with DNA binding dye DAPI. Traditional segmentation algorithms generally do not work on bright-field images and often work poorly on noisy signals such as those in immunofluorescence data. Therefore, our datasets serve as stress tests of the segmentation algorithm by presenting more challenging use cases.

We train modified UNets (Fig. S1) with the acquired images to generate a segmentation model. We first used the Cell Profiler software to analyze the nucleus image and computed the locations of each detected nucleus as the markers, which in turn were used to produce image patches (64×64 pixels), each centered around a marker location. The patch size was chosen to be larger than the size of a single cell, thus many patches have significant overlapping areas. The image patches were used as the inputs to train the neural network by minimizing the loss function *L*. The goal is to train the UNet to segment one single cell from each image patch. However, CNN output is dependent on local information only, and is not aware of the absolution positions of the pixel inputs. To ensure the segmentation apply to only one specific cell, we encoded positional information in the input data by including a image of a Gaussian disk (*σ* = 15) at the marker location as an additional input channel. On top of that, in order to ensure the features learned about cells are symmetric according to image orientation, we randomly flip the patches along two major axises. A corresponding reverse operation was performed on the network output before the computation of the loss function. For each independent dataset input, we optimize a separate segmentation model based on the same underlying network architect and training procedure.

The training process is stable and reasonably efficient. The loss function value (Fig. S2) decreases in a generally monotonic fashion until converging to a stable minimal value. The overall process took between 3-6 minutes on a single computational node with a K80 GPU (graphic processing unit), depending on the input. In addition, we quantified segmentation accuracies by computing mean intersection over union (mIOU) between machine segmentations and manual segmentations (more details below). We found that the segmentation accuracy increases monotonically over the training process and reached a stable maximum. Therefore, the training process did not exhibit overfitting defects, supporting the validity of the loss function design for the segmentation purpose. After the loss function value stabilized, we then integrated the single-patch outputs from the CNN to generate the segmentation map for the whole field-of-view (Fig. 2) for both immunofluorescence input and bright field input. A quick visual examination indicated that the algorithm yielded qualitatively believable segmentations in both cases.

**Figure 2.**
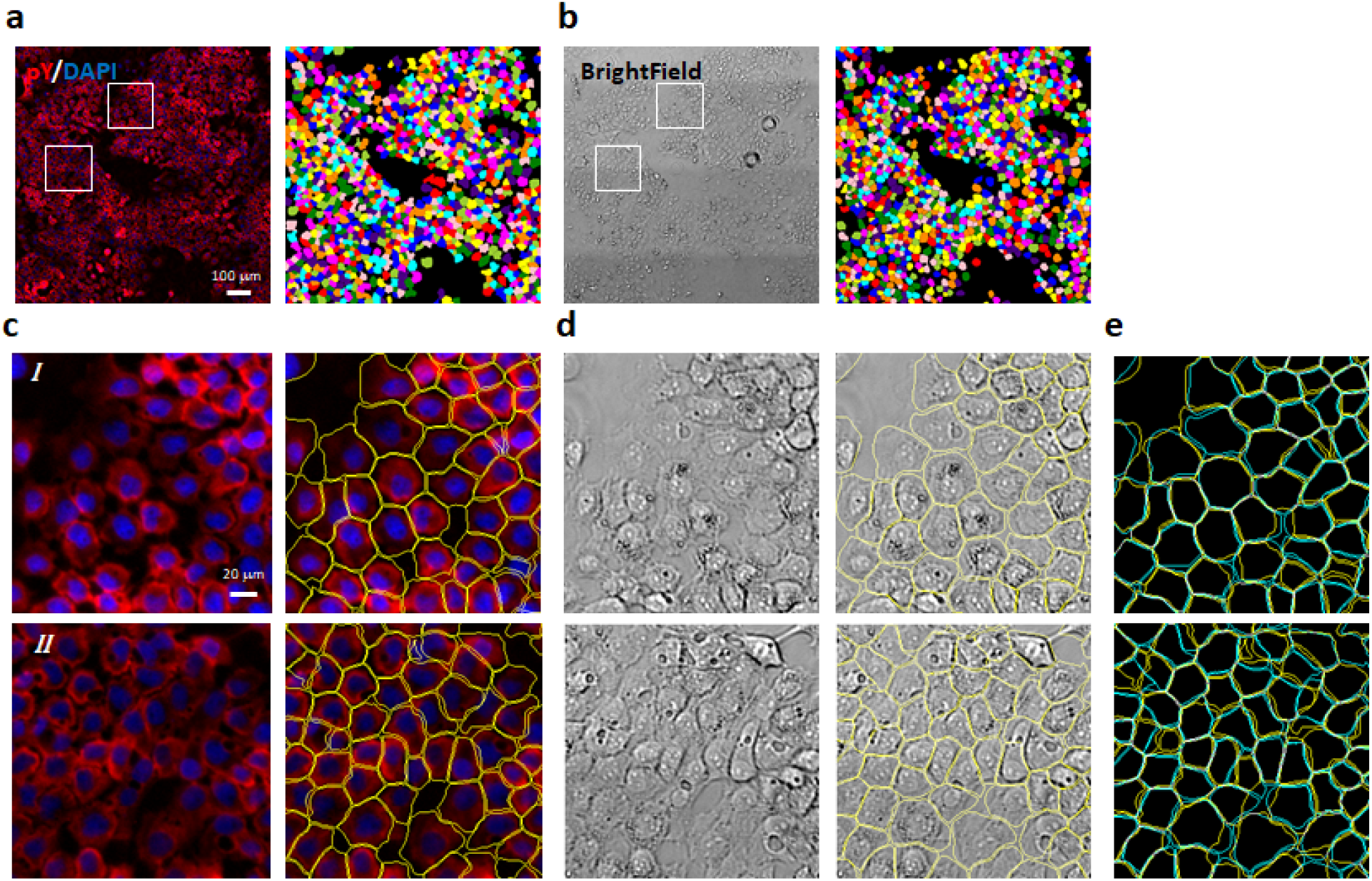
Segmentation of microscopy images by the CNN. Anti-pY immunofluorescence (**a**) and bright-field (**b**) images of A431 cells were acquired from the same sample location and the CNN segmentation results are shown as color maps (**a** and **b**, right panels). (**c** & **d**) Zoomed-in view of the segmentation results from the two subregions denoted by the white boxes in **a** & **b**. Here only the outline of the segmentations were plotted as image overlays (**c** & **d**, right columns). (**e**) Comparisons of the segmentation results based on the immunofluorescence images (yellow) versus the bright-field images (cyan). Data depict the same sub-regions as shown in **c** & **d**. All CNN segmentation models were trained with either the immunofluorescence image or the bright-field image only. DAPI images (shown in blue in **a** & **c**) were used for determining marker locations, but not used in training the CNN model.

**Figure 3.**
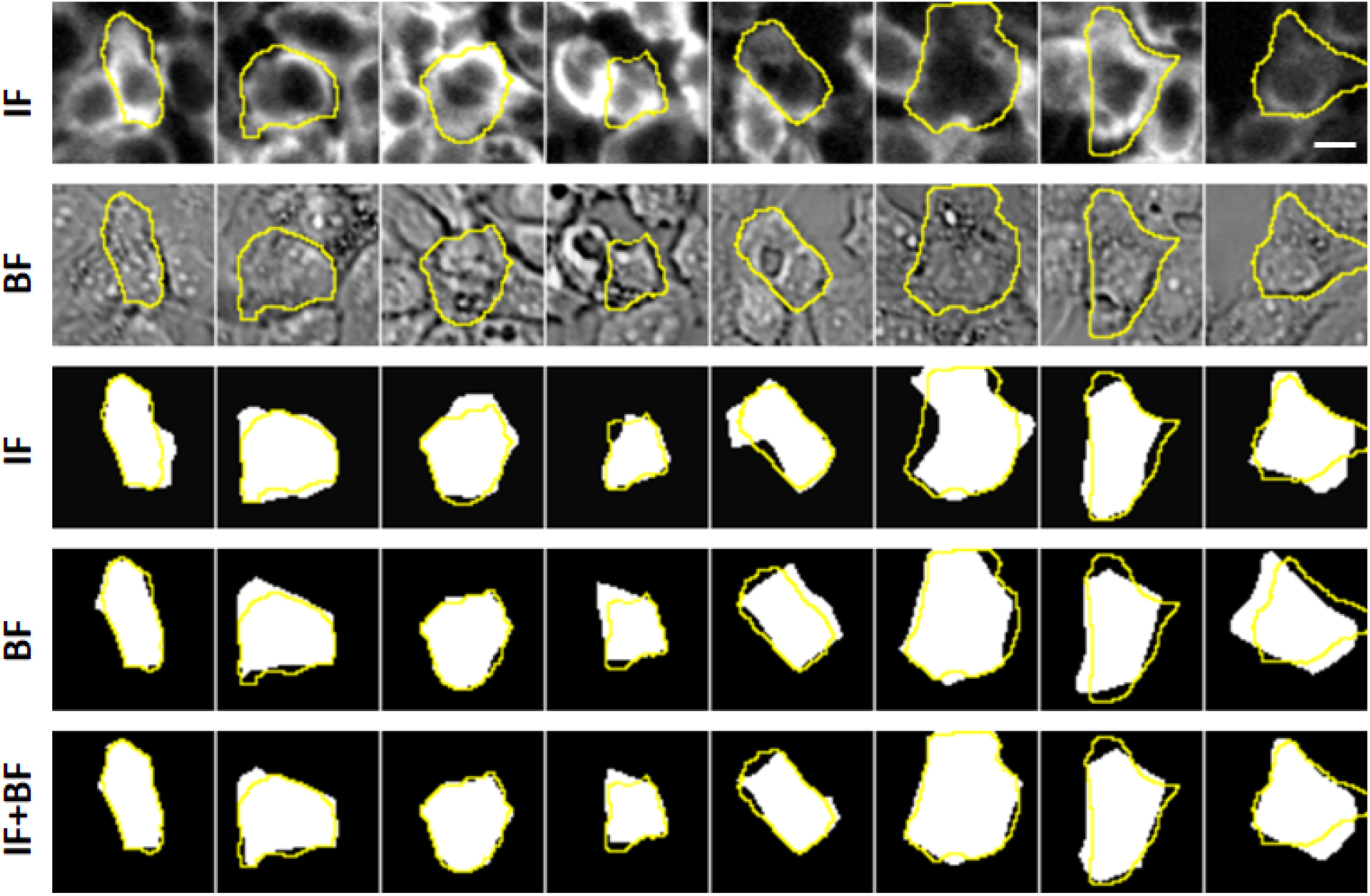
Comparisons between machine and manual segmentation results. Top two rows show the examples of individual image patches from the training data. The first row shows the patches from the anti-pY immunofluorescence (IF) images. The second row shows the patches from the bright-field (BF) images. The bottom three rows show the CNN segmentation output of each image patch. The models were trained with either the IF (row 3), BF (row 4) or multi-channel input of IF and BF combined (row 4). In all cases, the manual segmentations of the cells were plotted as the yellow overlays. Manual segmentation is based on a combined input of IF, BF and nucleus (DAPI) images (not shown here). The scale bar represents 20 μm.

To evaluate the segmentation quality, we performed three different quantitations of the segmentation results. First we quantified pixel level segmentation accuracy. To do that, we manually segmented ~130 cells randomly picked from three experimental datasets. The manual segmentation is performed by examining all available inputs (immuno-fluorescence, bright-field, and nucleus images) to ensure the highest accuracy. We then compute mIOUs between algorithm outputs and manual segmentations (Table I). As expected, the modal accuracy was higher when the model had taken multi-channel data as input during training. The highest mIOU accuracy (0.825) was achieved when the model was trained with all available data (immuno-fluorescence, bright-field, and nucleus images); conversely training the models with only the bright-field images resulted in the lowest accuracy (0.701). Training with immuno-fluorescence images produced better models than training with bright-field images, which is consistent with the impression that the immuno-fluorescence data offers clearer visual cues of cell boundaries. Including the nucleus images in the model training, somewhat surprisingly, improved the model accuracy slightly. Furthermore, inclusion of the nucleus images increased the convergence speed of the training significantly(Fig. S2). This is probably because the model could (correctly) infer that a segmentation boundary should traverse through the gap between two nuclei. Thus the exact information provided by the nucleus images allows quicker learning of the semantic features from the cell images. Finally, for a baseline analysis, we performed cell segmentation on the immunofluorescence data using Cell Profiler, a well-known cell segmentation software (Table 1). Cell profiler offers several predefined algorithms for unsupervised segmentation, the best of which resulted in an mIOU value of 0.69, significantly worse than our CNN model on the same fluorescence data. Cell Profiler was not designed to perform segmentation on bright-field images, therefore no comparison was made for the segmentations of bright-field image data.

**Table I.**
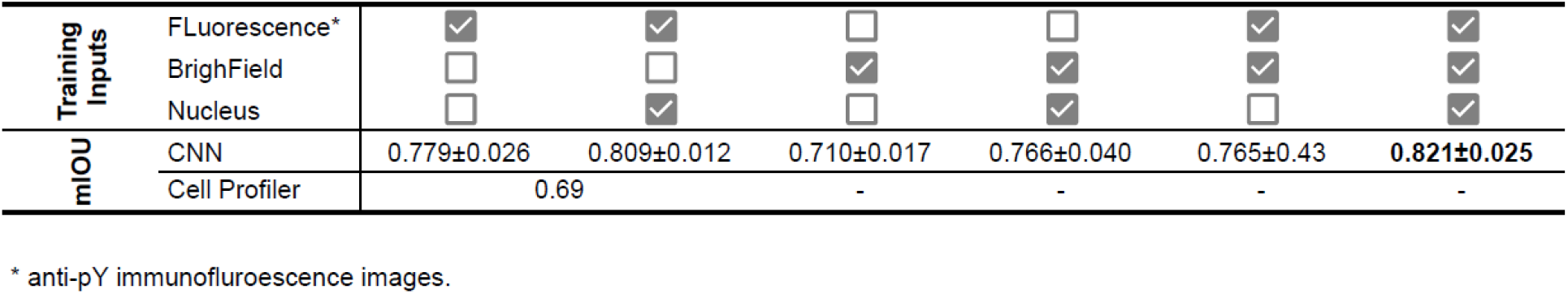
Unsupervised CNN segmentation accuracy.

Next we evaluated the consistency of segmentation results on images of differential modalities. One advantage of our algorithm is that it does not assume a priori specific characteristics of the data, and instead try to “learn” the semantic features from the images themselves. As a result the same algorithm could perform segmentation on either the fluorescence images or the bright-field images (Fig. 2) without the need to change parameters. However, if the segmentation results were accurate, then the results should be similar, if not identical, on images representing the same sample, which we found to be the case (Fig. 2e).. Quantitatively, we found that the mIOU between two segmentation results, one on the anti-pY immunofluorescence images and the other on bright-field images, was 0.74±0.017. The mIOU value increased to 0.809±0.051, if the nucleus images were also included during the model training. Therefore the segmentation outputs on differential image modalities indeed yielded results similar to one other, further supporting the notion that the self-supervised training process allows the CNN to learn correct cellular features.

Additionally, we tested whether models trained with one image input can perform segmentation on a different image that it had not seen yet. If the unsupervised training procedure allows the model to learn semantic features, then we should reasonably expect the trained model to also be able to operate on a different image that is acquired in a similar manner. Indeed we found that a pre-trained model can segment an unseen image producing a segmentation similar to that of a “proper” model, which had previously seen the image (Fig. S3), with mIOU = 0.752 for anti-pY immunofluorescence images and mIOU = 0.772 for bright-field images.

Next, we explore the possibility of generating marker positions directly from the cellular images themselves (i.e, not requiring a nucleus image). While the existence of the nucleus image provides a convenient means to generate the marker locations, the requirement exerts extra experimental overhead. We hypothesized that marker locations can also be computationally obtained, if we produce a CNN model that could produce a synthetic “nucleus” image from a normal cellular image, e.g., a bright-field image of cells. We tested this idea by first training a UNet to perform a mapping, either from the anti-pY immunofluorescence images or from the bright-field images, to their respective nucleus images. It is important to note that our goal is not to produce a realistic nucleus image, but to compute the mark locations. Thus the model can be of relatively low resolution. Indeed, we found that a useful model can be obtained with very little training data: i.e., two images of 1750×1750 pixels in our test. Fig. 4 shows examples of synthetic nucleus images by applying pretrained models to new images the models hadn’t seen. While these synthetic images were far from indistinguishable from real nucleus images, they reproduced the nucleus positions with enough accuracy to allow computing marker locations via a simple blob detection algorithm. The marker locations were then combined with the original cellular images to train a segmentation model using the unsupervised procedure outlined earlier.

**Figure 4.**
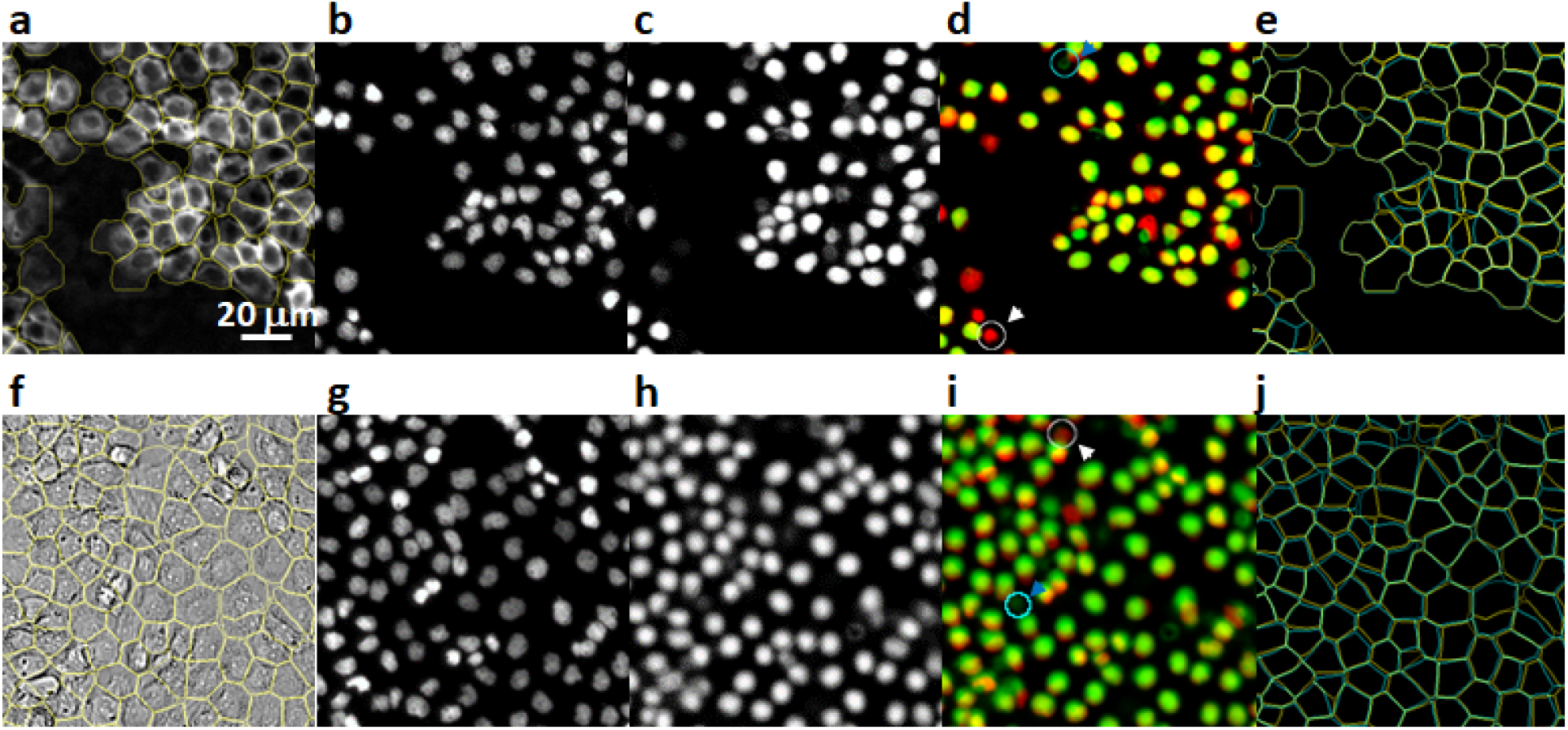
Segmentation based on synthetic marker locations. Results for the anti-pY immunofluorescence data are shown in panel **a**-**e**, and results for the bright-field image are shown in(**f**-**j)**. The input images (**a**& **f**) were shown together with the segmentation results (yellow overlay). Experimentally acquired nucleus (DAPI) images were shown in **b**& **g** for comparison, although they were not used for the computation of the segmentation. Instead, synthetic nucleus images (**c**& **h**) were computed directly from the input images (**a**& **f**) using a pre-trained CNN. Errors in the synthetic images can be easier discerned in the composite overlay images (**d**& **i**) of the experimental (red) and synthetic (green) nucleus images. Both false negative (white arrow) and false positive (blue arrow) errors were found, but are of relatively low occurrences. Finally, the segmentation results based on synthetic markers were compared with the “proper” segmentation, for which the experimental DAPI images were used for marker locations. The comparisons were shown in **e**& **j**, where segmentations based on experimental markers (yellow) were drawn in overlay on segmentations based on synthetic markers (cyan).

To evaluate the accuracy of the segmentation based on this synthetic marker procedure, we separately quantify two sources of errors. Firstly, the model generated synthetic markers did not perfectly match each individual nucleus in the sample, which resulted in either missing markers (false negative) or extra markers (false positive) at wrong locations (Fig. 4). We compared the synthetic marker locations with experimental images of the nuclei and found that, for markers generated from immunofluorescence images, the average false negative rate is 1.8±0.3% and the average false positive rate os 2.2±0.3%, and for markers generated from bright-field images, the average false negative rate is 2.1±0.2% and the average false positive rate os 2.4±0.3%. These two types of errors correspond to missing cells and over-segmentation of cells, respectively, in the final segmentation results. Furthermore, the segmentation model had additional segmentation error at pixel level, which we evaluated by comparing the single cell segmentation results with manual segmentations on selected cells. We found the mIOU value to be 0.814±0.025 for segmenting anti-pY immunofluorescence images and 0.755±0.033 for segmenting bright-field images. Interestingly, the segmentation errors produced here are not significantly different from the models trained with “correct” marker locations (i.e, using experimental nucleus images, see Table 1), indicating that the small amount of false-positive and false-negative errors in synthetic markers had little impact on the training of the segmentation models.

## Discussion

Cellular imaging based high content screening has become more widely used in biological and medical researches^23,24^. The ability to perform single cell segmentation accurately and in a cost-effective manner is of great importance both to the commercial interests and to the basic sciences of studying single cells. Currently, accurate segmentation often requires a tailored algorithm specific to a fixed imaging modality, labeling procedure and specific imaging settings. Deviation from an optimized protocol often resulted in lowered performances. In general, these requirements significantly limit the flexibility in experimental designs. Machine learning based methods, on the other hand, promise to be more flexible because they are able to learn correct semantic features directly from data. Indeed, we show here that a CNN based computation method can perform cell segmentation on inputs of vastly different signal characteristics. In comparison to conventional algorithm-based segmentation tools, our method has the advantage of having a higher accuracy and being applicable to various input signal modality without the need of parameter tuning. More importantly, unlike previous CNN based segmentation tools, our method does not require any human labeled “ground-truth” data for training, and instead rely on an unsupervised training procedure, alleviating a significant barrier in applying deep learning strategy to cell segmentation.

The key insight from our work is that a properly designed artificial intelligence model can learn semetic features from unlabeled cell image data by processing and comparing multiple small image patches. The general concept of patch learning was recently introduced by several highly influential studies^25,26^ of unsupervised machine learning. Most current works in this area, however, are focusing on the spatial context between non-overlapping patches as a source of free information for training the models. The training generally requires a large amount of input data. On the other hand, cellular images produced in biology have significantly higher feature density (i.e., they contain nothing but cells) than most natural images. We show that the unique characteristics of cell images can be leveraged to reduce the training data required, to a single input image in our case. Tailored to this task, our training method used overlapping image patches and relied on the consistency between patches for training, resulting in a unique algorithm for this task.

The accuracy of our method is the highest if the user also acquires nucleus images from the sample, which allows computer generation of “markers” that signify the location of individual cells. However, the experimental data is optional, as we show “synthetic” nucleus images can be generated using a machine learning approach. The latter strategy does require pre-training of a CNN, which in turn requires additional training data in the form of cellular image / nucleus image pairs. We note that nominally this is a type of supervised learning, but the training process does not require manually labeling of data, and uses experimentally acquired images as the “ground truth” instead. The overhead involved in the training process is generally low -- in our case, we showed that training with two pairs of images were sufficient to produce results accurate enough for marker computation. However, the training is specific to the exact signal of interest. Switching the cellular image to a different label would need retraining of the network. In the end, users of the algorithm will decide which strategy (experimental vs synthetic nucleus markers) is more compatible with the specific research goal at hand.

## Method

### Cell culture and microscopy

All experiments were performed using the human squamous-cell carcinoma line A431. Cells were plated onto plasma cleaned cover glass and grown to ~70% confluence in standard growth media. Cells were quickly washed once with PBS and fixed with 4% paraformaldehyde for 20 min. Cells were then washed with PBS three times for 5 min each, permeabilized in 1.5% Triton-X PBS for 10 min and washed with PBS three times for 10 min each. Dishes were then blocked in 1.5% BSA PBS for 60 min at 4°C, incubated with alexa647-labeled anti-pY (1:500 in blocking buffer) for 2 h while rocking at 4°C, washed three times for 5 min in PBS, and imaged in PBS in presence of DAPI. Epi-fluorescence and brightfield images were acquired on an inverted fluorescence microscope (Olympus IX73) with an 20x objective. An area of 1.4×1.4 mm was imaged with a grid pattern for each sample, resulting in stitched images of 1750×1750 pixels. The final images were stitched using the stitching plugin available in the Fiji software. Datasets were collected from three sample replicates.

### Model training

We first obtain a list of marker positions according to the centroid locations of nuclei, using Cell Profiler (for experimental nucleus images) or using a blob detection algorithm (for synthetic nucleus images). Image patches were extracted from the inputs, which were either anti-pY fluorescence images or bright-field images of cells. A modified UNet, used as the engine for cell segmentation, was trained to minimize the custom loss function (Eq 1) with fixed hyperparameter values of λ = 1 and β = 15, using the adaptive momentum estimation (ADAM) optimization algorithm. Typically, processing all available image patches in one batch would exceed the memory capacity of a GPU, even if the patches are from just one input image. Therefore, we limit each computational batch to patches within a 640×640 area of the input image and process the whole image in steps. Data augmentation was implemented by randomly flipping and/or transposing the input patches. A corresponding reverse operation was performed on the network output before the computation of the loss function. The network was trained until the loss function reaches a stable minimum. To integrate individual segmentation results from patches to yield the segmentation results for the whole field-of-view, each pixel was assigned to the cell with the highest probability output at that location. Pixels showing segmentation probabilities less than 0.5 from all patches were assigned as the background.

### Binary mask

A binary mask was used during training to ensure that the segmentation of individual cells will not exceed the area covered by the cells. For fluorescence images, the binary mask is generated using the “Graph Cut”^27^ plugin in the Fiji software, which implements the well-known graph-cut algorithm. The algorithm relies primarily on the gray value differences to separate the foreground from the background, but compensate for the over-segmentation by introducing a penalty for foreground/background boundaries. For bright-field images, the gray values are not useful for distinguishing the foreground from the background. Therefore, we first train a two-class RandomForest model using the “Weka Segmentation”^28^ Fiji plugin. We used Gaussian filter, variance and laplacian values as feature inputs. We then performed a graph cut on the model’s probability output and used the results as the mask.

### Synthetic nucleus image

The same UNet (Fig. S1) architecture was used as the base CNN model for producing synthetic nucleus images. The UNet models were trained with small datasets of either two immunofluorescence (anti-pY) images or two brightfield images. Each input image was paired with a nucleus image, which was thresholded and converted to binary images. The training was stopped when the cross-entropy loss between model output and the (binary) nucleus images reached the value of 0.15. The pre-trained model was applied to new input images to produce the synthetic nucleus images.

## Code availability

Current implementation of the segmentation algorithm is available as a github repository. https://github.com/jiyuuchc/cellcutter

---

**Figure S1.**
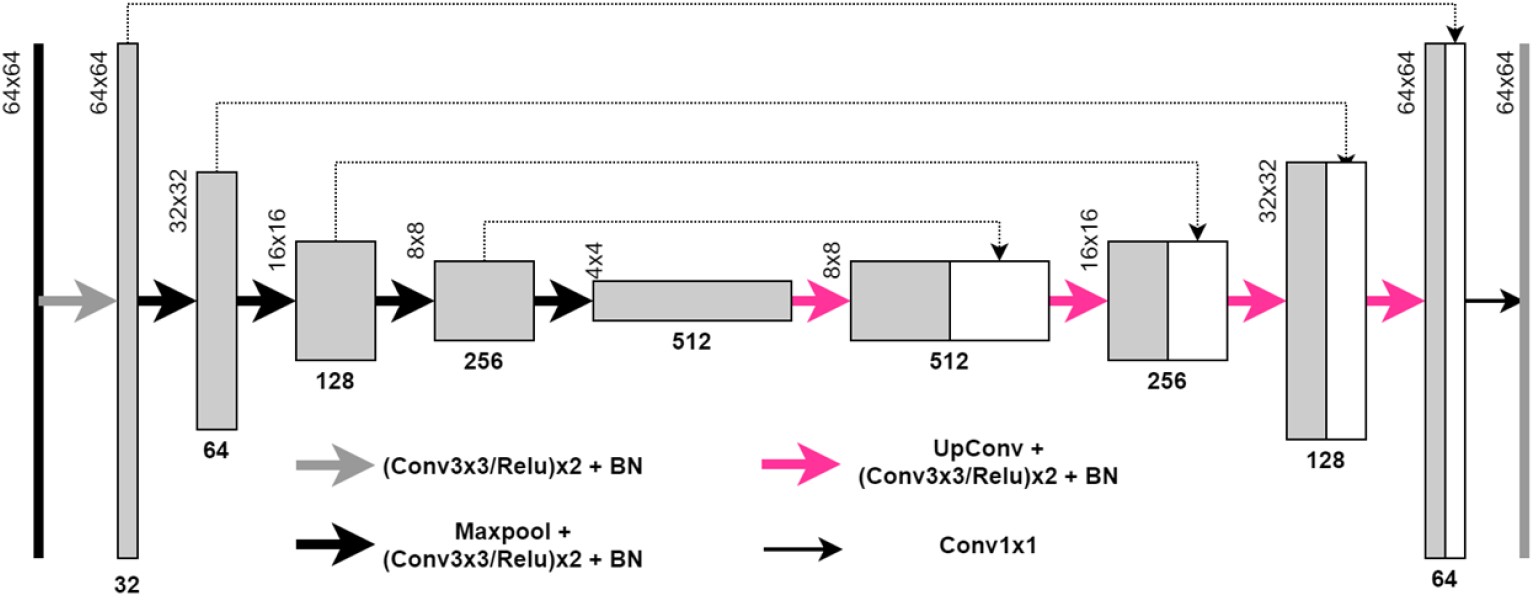
Architecture of the CNN used for segmentation. The encoder portion of the network extracts image features by multiple stages of convolution and max pooling. The decoder portion of the network constructs the segmentation by incremental upscaling of the feature map (to 2x larger dimension) followed by convolution. Feature map obtained in the decoding step is concatenated by intermediate results at the decoding steps in order to supplement feature information at incrementally higher resolution. Relu activation is used throughout the encoding and decoding steps, except in the final output layer, in which a sigmoid activation is used. The same network is used for producing synthetic nucleus images, in which case the input tensor dimension is much larger (870×870) to match the size of training data.

**Figure S2.**
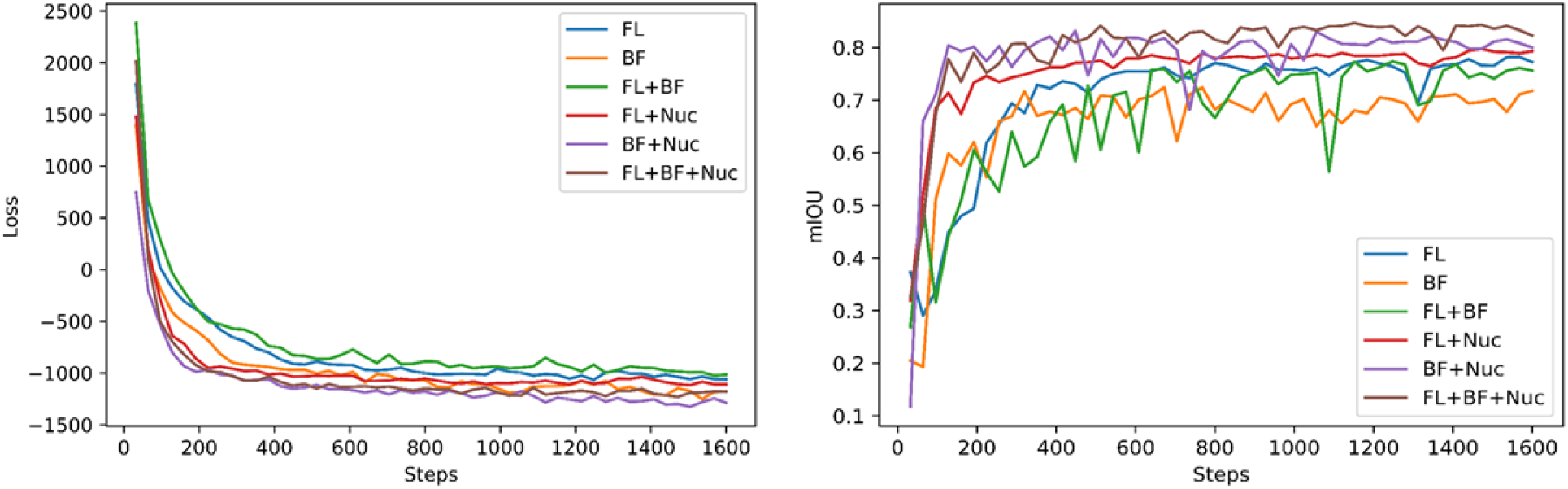
Representative training traces showing the minimization of the loss function (left) and the corresponding optimization of the segmentation accuracies (right) over the self-supervised training process. Multiple models were trained with various combinations of input data, including the immunofluorescence image (FL), the bright-field image (BF) and the DAPI staining image (Nuc).

**Figure S3.**
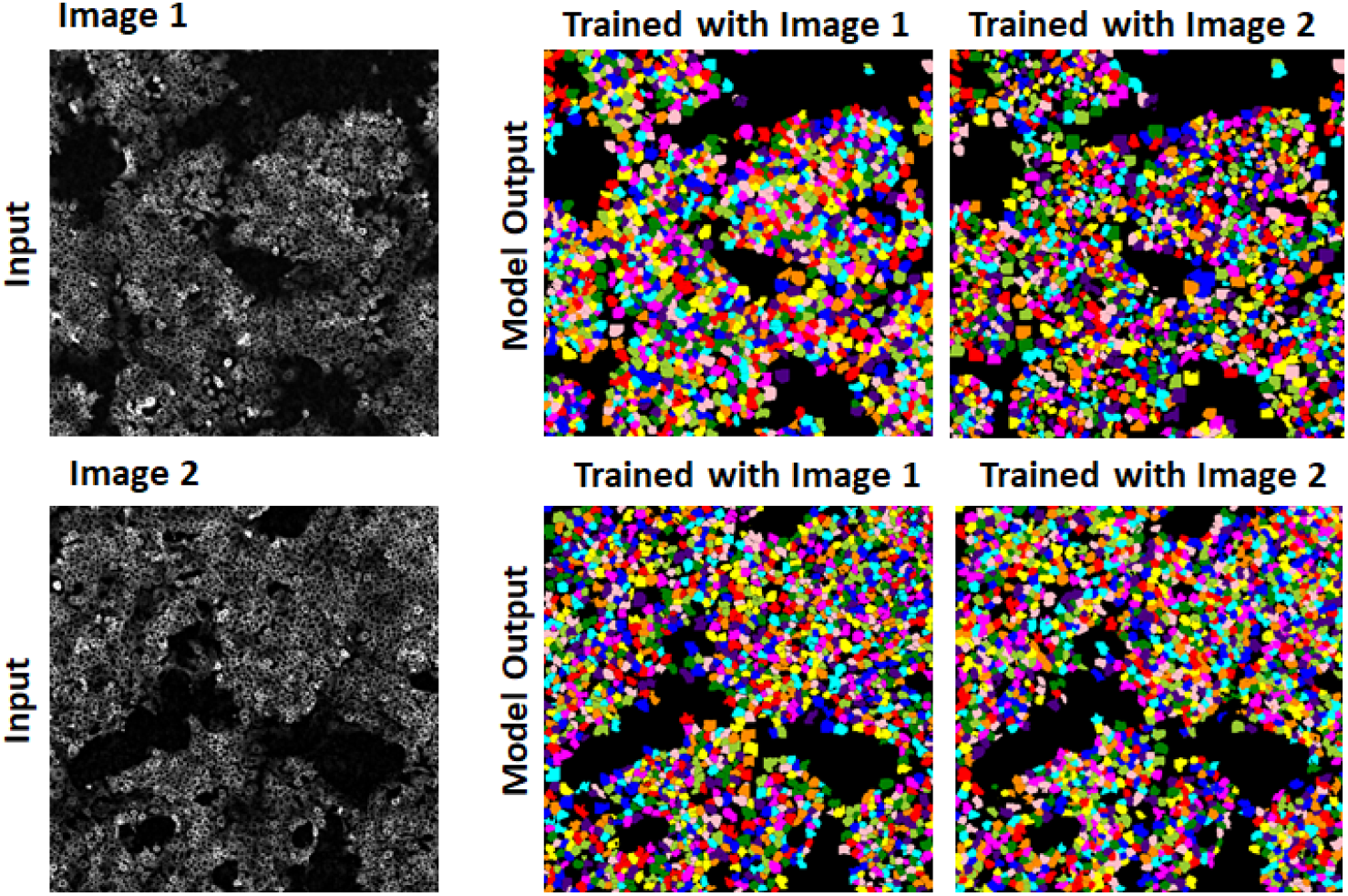
Comparison of segmentation results on both seen and unseen fluorescence images. Two segmentation models were separately trained by two immunofluorescence images of the same modality (left) and the resulting models were used to operate on both images (one seen and one unseen) to obtain the segmentation (right). The results demonstrated that the models trained this way were able to apply learnings to unseen data of similar features.

**Figure S4.**
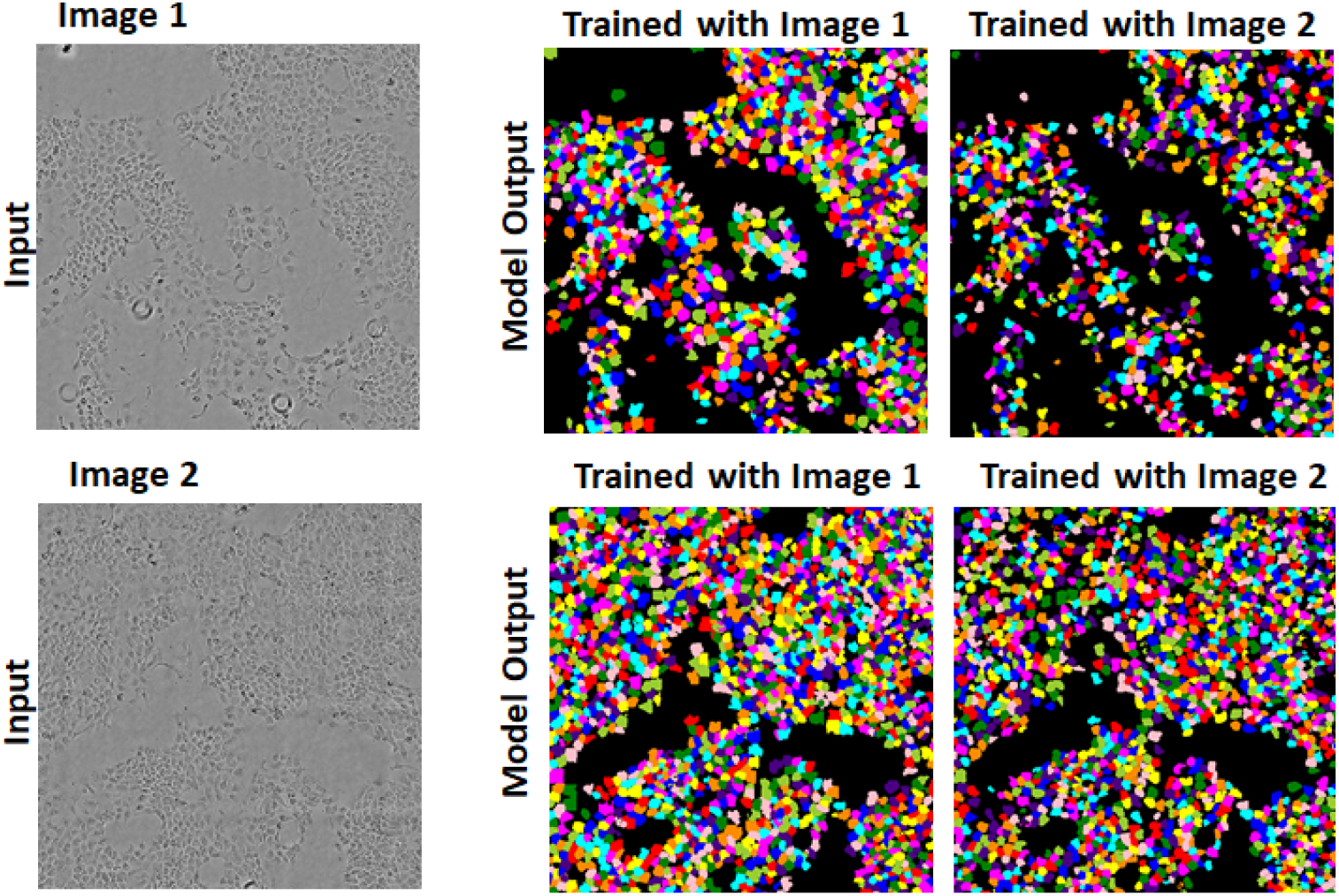
Comparison of segmentation results on both seen and unseen bright-field images. This is the same analysis as in Fig S3 except on bright-field images.

## References

1. Meijering, E. Cell Segmentation: 50 Years Down the Road [Life Sciences]. IEEE Signal Processing Magazine 29, 140–145 (2012).

2. Deshmukh, B. S. & Mankar, V. H. Segmentation of Microscopic Images: A Survey. in 2014 International Conference on Electronic Systems, Signal Processing and Computing Technologies 362–364 (2014). doi:10.1109/ICESC.2014.68.

3. Krizhevsky, A., Sutskever, I. & Hinton, G. E. ImageNet classification with deep convolutional neural networks. Commun. ACM 60, 84–90 (2017).

4. Long, J., Shelhamer, E. & Darrell, T. Fully convolutional networks for semantic segmentation. in 2015 IEEE Conference on Computer Vision and Pattern Recognition (CVPR) 3431–3440 (2015). doi:10.1109/CVPR.2015.7298965.

5. Ronneberger, O., Fischer, P. & Brox, T. U-Net: Convolutional Networks for Biomedical Image Segmentation. arXiv:1505.04597 [cs] (2015).

6. Liu, Y., Minh Nguyen, D., Deligiannis, N., Ding, W. & Munteanu, A. Hourglass-ShapeNetwork Based Semantic Segmentation for High Resolution Aerial Imagery. Remote Sensing 9, 522 (2017).

7. Al-Kofahi, Y., Zaltsman, A., Graves, R., Marshall, W. & Rusu, M. A deep learning-based algorithm for 2-D cell segmentation in microscopy images. BMC Bioinformatics 19, 365 (2018).

8. Cameron, W. D., Bennett, A. M., Bui, C. V., Chang, H. H. & Rocheleau, J. V. Leveraging multimodal microscopy to optimize deep learning models for cell segmentation. APL Bioengineering 5, 016101 (2021).

9. Wang, W. et al. Learn to segment single cells with deep distance estimator and deep cell detector. Computers in Biology and Medicine 108, 133–141 (2019).

10. Wollmann, T. et al. GRUU-Net: Integrated convolutional and gated recurrent neural network for cell segmentation. Medical Image Analysis 56, 68–79 (2019).

11. Wolny, A. et al. Accurate and versatile 3D segmentation of plant tissues at cellular resolution. eLife 9, e57613 (2020).

12. Bertrand, G. On Topological Watersheds. J Math Imaging Vis 22, 217–230 (2005).

13. Carpenter, A. E. et al. CellProfiler: image analysis software for identifying and quantifying cell phenotypes. Genome Biology 7, R100 (2006).

14. Berg, S. et al. ilastik: interactive machine learning for (bio)image analysis. Nature Methods 16, 1226–1232 (2019).

15. Wang, M. et al. Novel cell segmentation and online SVM for cell cycle phase identification in automated microscopy. Bioinformatics 24, 94–101 (2008).

16. Sharif, J. M., Miswan, M. F., Ngadi, M. A., Salam, M. S. H. & bin Abdul Jamil, M. M. Red blood cell segmentation using masking and watershed algorithm: A preliminary study. in 2012 International Conference on Biomedical Engineering (ICoBE) 258–262 (2012). doi:10.1109/ICoBE.2012.6179016.

17. Ulman, V. et al. An objective comparison of cell-tracking algorithms. Nature Methods 14, 1141–1152 (2017).

18. Dzyubachyk, O., Niessen, W. & Meijering, E. Advanced level-set based multiple-cell segmentation and tracking in time-lapse fluorescence microscopy images. in 2008 5th IEEE International Symposium on Biomedical Imaging: From Nano to Macro 185–188 (2008). doi:10.1109/ISBI.2008.4540963.

19. Al-Kofahi, Y., Lassoued, W., Lee, W. & Roysam, B. Improved Automatic Detection and Segmentation of Cell Nuclei in Histopathology Images. IEEE Transactions on Biomedical Engineering 57, 841–852 (2010).

20. Ounkomol, C., Seshamani, S., Maleckar, M. M., Collman, F. & Johnson, G. R. Label-free prediction of three-dimensional fluorescence images from transmitted-light microscopy. Nature Methods 15, 917–920 (2018).

21. Kingma, D. P. & Ba, J. Adam: A Method for Stochastic Optimization. arXiv:1412.6980 [cs] (2017).

22. Martín Abadi et al. TensorFlow: Large-Scale Machine Learning on Heterogeneous Systems. (2015).

23. Boutros, M., Heigwer, F. & Laufer, C. Microscopy-Based High-Content Screening. Cell 163, 1314–1325 (2015).

24. Zanella, F., Lorens, J. B. & Link, W. High content screening: seeing is believing. Trends in Biotechnology 28, 237–245 (2010).

25. Doersch, C., Gupta, A. & Efros, A. A. Unsupervised Visual Representation Learning by Context Prediction. in 1422–1430 (2015).

26. Noroozi, M. & Favaro, P. Unsupervised Learning of Visual Representations by Solving Jigsaw Puzzles. in Computer Vision – ECCV 2016 (eds. Leibe, B., Matas, J., Sebe, N. & Welling, M.) 69–84 (Springer International Publishing, 2016). doi:10.1007/978-3-319-46466-4_5.

27. Boykov, Y. & Kolmogorov, V. An experimental comparison of min-cut/max-flow algorithms for energy minimization in vision. IEEE Transactions on Pattern Analysis and Machine Intelligence 26, 1124–1137 (2004).

28. Arganda-Carreras, I. et al. Trainable Weka Segmentation: a machine learning tool for microscopy pixel classification. Bioinformatics 33, 2424–2426 (2017).

